# Assessment of ecological fidelity of human microbiome-associated mice in observational studies and an interventional trial

**DOI:** 10.1101/2025.03.11.642547

**Authors:** Matthew K. Wong, Eric Armstrong, Alya A. Heirali, Pierre H.H. Schneeberger, Helen Chen, Kyla Cochrane, Keith Sherriff, Emma Allen-Vercoe, Lillian L. Siu, Anna Spreafico, Bryan Coburn

## Abstract

Composition and function of the gut microbiome is associated with diverse health conditions and treatment responses. Human microbiota-associated (HMA) mouse models are used to establish causal links for these associations but have important limitations. We assessed the fidelity of HMA mouse models to recapitulate ecological responses to a microbial consortium using stools collected from a human clinical trial. HMA mice were generated using different routes of consortium exposure and their ecological features were compared to human donors by metagenomic sequencing. HMA mice were more similar in gut composition to other mice than their respective human donors, with taxa including *Akkermansia muciniphila* and *Bacteroides* species enriched in mouse recipients. A limited repertoire of microbes was able to engraft into HMA mice regardless of route of consortium exposure. In publicly available HMA mouse datasets from four distinct health conditions, we confirmed our observation that a taxonomically restricted set of microbes reproducibly engrafts in HMA mice and observed that stool microbiome composition of HMA mice were more like other mice than their human donor. Our data suggest that HMA mice are limited models to assess the ecological impact of microbial consortia, with ecological effects in HMA mice being more strongly associated with host species than donor stool ecology or ecological responses to treatment in humans. Comparisons to published studies suggest this may be due to comparatively large host-species effects that overwhelm ecological effects of treatment in humans that HMA models aim to recapitulate.

**Importance:** Human microbiota-associated (HMA) mice are models that better represent human gut ecology compared to conventional laboratory mice and are commonly used to test the effect of the gut microbiome on disease or treatment response. We evaluated the fidelity of using HMA mice as avatars of ecological response to a human microbial consortium, MET4. Our results show that HMA mice in our cohort and across other published studies are more similar to each other than the human donors or inoculum they are derived from and harbour a taxonomically restricted gut microbiome. These findings highlight the limitations of HMA mice in evaluating the ecological effects of complex human microbiome-targeting interventions, such as microbial consortia.

## Introduction

Many studies have identified associations between gut microbial community composition and pathological or physiological phenotypes. These discoveries have led to an increasing number of human interventional trials targeting the microbiota in a broad range of health conditions. Germ-free and gnotobiotic mice have been instrumental in exploring these interactions. Human microbiota-associated (HMA) mice are generated by fecal microbiota transplant (FMT) of human donor stool into germ-free or antibiotic-treated mice to recreate human donor ecology and assess associated pathology and other phenotypes. This approach provides a model by which phenotypic differences can be causally attributed to compositional differences in or between the human donors’ gut microbiomes and provides a platform for interrogating mechanism with experimental strategies that would be impractical or impossible in humans.

Although HMA mouse models have become a common tool for inferring causal effects of the gut microbiome in clinical studies, they have been subject to criticism and scrutiny(1–4). Differences in species physiology, diet, and immunity are pivotal in determining which microbes can establish a niche in the murine gut. Thus, although these models have demonstrated utility as reductionist models where the inoculum is defined and limited, more complex microbial communities may only partially recapitulate donor ecology. As such, care must be taken when designing and interpreting results from HMA mouse models generated with diverse and complex communities.

Microbial consortia are curated communities of microbes that are designed to be a safer and more reproducible alternative to fecal microbiota transplant(5). Therapeutic mechanisms of disease modification by consortia have been assessed in mouse models of a diversity of conditions in which the microbiome has been causally implicated, such as *Clostridioides difficile* infection(6) and non-infectious conditions such as cancer(7, 8). A recent clinical trial, MET4-IO(9), tested for the safety and engraftment of Microbial Ecosystem Therapeutic 4 (MET4), a microbial consortium designed for combination with cancer immunotherapy. This presented an opportunity to study the ecological fidelity of HMA mouse models as ‘avatars’ of human ecological responses to a microbiome-targeting intervention.

Here, we generated HMA mice using stool from donors that participated in MET4-IO with the purpose of assessing the concordance between ecological responses to a microbial consortium between HMA mice and their corresponding donors. Ecological recapitulation in HMA mouse recipients of human microbiomes from published datasets was then performed to determine how generalizable our results were. The goals of this study were to assess how well HMA mice represented human ecological changes that occur during therapeutic administration of microbial consortia and ascertain their suitability as experimental avatars in this context.

## Methods

### Study cohort and sample collection

Individuals with solid tumours were enrolled into an early phase clinical study that assessed the safety, tolerability, and engraftment of MET4, a novel microbial consortium(9). Participants from “cohort B” were the primary focus of this study, consisting of immune checkpoint inhibitor (ICI)-naïve participants who were stratified into MET4 + standard-of-care ICI (anti-PD-1 alone, or combination with anti-CTLA-4) or ICI alone (no MET4) treatment groups. Further details of the trial are detailed in the MET4-IO trial publication(9).

Paired participant stool samples were collected from MET4-IO cohort B participants from both MET4-treated and control groups. Samples were collected 3-4 weeks after initiation of ICI but prior to MET4 (T0), and 3-4 weeks after initiation of MET4, while on ICI (window +2 weeks, T2). If a T2 sample was not available, an end of therapy (EOT) sample was used if it was collected no more than 2 weeks after the scheduled timing for T2. Stools were frozen at −80^°^C.

### MET4 consortium production and composition

MET4 consists of 30 individually cultured bacterial isolates chosen based on association with ICI response(10–12). Bacterial strains were encapsulated together and assessed for consistency between drug batches. The taxonomic composition of MET4 has previously been reported(9) and is included in Supp. Table 1 along with taxonomic annotations used in our analyses.

**Table 1.** Studies included in HMA mouse mega-analysis.

| Study | Condition | Sequencing type | # of donor-mouse groups used |
| --- | --- | --- | --- |
| Britton et al.(20) | Intestinal bowel disease | 16S | 15 |
| Sampson et al.(21) | Parkinson's disease | 16S | 12 |
| Baxter et al.(22) | Colorectal cancer | 16S | 6 |
| Sharon et al.(23) | Autism spectrum disorder | 16S | 8 |

### Stool processing

PBS + 20% glycerol was degassed in an anaerobic chamber overnight prior to stool processing. Stools were thawed and resuspended at 10mL/g in degassed PBS + 20% glycerol to make a fecal slurry. Slurries were passed through a 300 µm Whirl-Pak filter bag (Nasco) under anaerobic conditions to remove stool particulates. Filtered fecal slurries were frozen at −80^°^C until use.

### Human microbiota-associated mice

To generate the HMA mice, germ-free mice were gavaged with 250µL of thawed human donor fecal slurries. Extra slurry was lathered onto animal fur and cage bedding. Cages were kept in germ-free conditions for two weeks before being moved to specific pathogen-free (SPF) housing conditions, where mice were not handled for a week to allow the gut microbiome to stabilize. Stools were collected at baseline, one-week post-SPF entry, and 3-4 weeks after SPF entry.

Using this method, paired pre/post-MET4-treated HMA mice were generated with two ‘routes’ of MET4 exposure. For the first, which we termed the ‘human-treated route’, mice were generated using paired stool samples obtained at T0 and T2 from 6 study participants (participants B001, B002, B004 and B005, B012 and B018), i.e. before and after MET4 administration to those participants. For the alternative method, which we termed the ‘mouse-treated route’, HMA mice were generated using T0 (pre-MET4 treatment) stools from participants B004 and B005 (both with increases of >5 MET4 taxa by >10-fold after treatment). Once the gut microbiome was stabilized, mice were treated directly with MET4. One week after SPF entry, MET4 was processed and diluted in 5 mL of degassed PBS anaerobically. 250µL of MET4 was gavaged into mice for three consecutive days and every 3 days for 2 weeks thereafter. Stools were collected at baseline, SPF entry, and 2 weeks post-MET4 initiation.

### Metagenomic sequencing

DNA was extracted from donor fecal slurries and mouse stools using Qiagen’s DNeasy Powersoil Pro kit. Illumina’s DNA Prep kit was used to prepare libraries from these samples. Nextera 96-well CD Indexes (Illumina) were used to index each sample before sequencing on an Illumina MiniSeq with a MiniSeq High Output Reagent Kit (Illumina) to obtain approximately 500,000 reads/sample (25 million total reads).

### Sequence data processing

Sequence quality was assessed with FastQC v.0.11.9(13). As the quality was high, no sequence trimming was performed. Nextera adapters were trimmed with Trimmomatic v0.39(14). Human, phiX, and mouse reads were removed with KneadData v.0.7.2(15). Taxa were identified from cleaned reads using Metaphlan v.4.0.6 using default settings(16, 17).

### Microbiome statistical analysis

MET4 taxonomic annotations used for analyses are listed in Supp. Table 1. Relative abundance (RA) % was compared between stools collected from mice generated from each donor-mouse pair at T0 and T2 timepoints. RA % were averages for mice generated with a single donor stool to avoid pseudo-replication(1, 2). Bray-Curtis dissimilarity matrices were generated with the “vegdist” function in the *vegan* package in R(18), using species-level taxonomy tables as input. Pairwise dissimilarities were manually extracted from the resulting dissimilarity matrix for group-wise comparisons. For MaAsLin2 analysis, taxonomic tables were filtered by removing species that contributed less than 0.1% relative abundance in one sample. The MaAsLin2 package v1.7.3(19) was used to identify taxa that were enriched in humans versus mice. Fixed effect was host species (i.e., human vs mouse). For models that included repeated measurements from individual participants, participant ID was included as a random effect. Default MaAsLin2 analysis parameters were used, and false discovery rate was used to control for multiple comparisons. A *q* value of 0.05 was considered significant.

### Mega-analysis on human microbiota-associated mouse studies

Four HMA mouse studies across distinct health states (inflammatory bowel disease, Parkinson’s disease, colorectal cancer, and autism spectrum disorder) were selected for a mega-analysis (20–23). Studies were selected based on public availability of raw 16S rRNA sequencing data for human donors and mice and appropriate study design (i.e. FMT of human donor stool from to recipient mice). Raw sequencing data were analyzed with QIIME2 v2024.2(24). Quality filtering and denoising was performed with DADA2(25). Taxonomic assignment of sequences was performed using a naïve Bayes classifier trained on the SILVA 132 99% operational taxonomic unit database(26, 27). Amplicon sequence variant (ASV) tables were defined at the genus level.

For Pearson correlation coefficient comparisons, a Fisher z-transformation was performed on coefficients prior to analysis. Percent engraftment was calculated for each taxon by dividing total human-mouse pairs with the taxon present in both by total human donors with the taxon present overall. For engraftment plots, only taxa engrafting in >4 mice were included. Bray-Curtis dissimilarity was generated and analyzed as above.

## Results

### Post-treatment stool composition differs in human donors and mouse recipients

We first sought to assess the reproducibility of human ecology in HMA mice generated using six sets of paired stools collected pre- (on ICI, before MET4) and post-MET4 (on ICI and on MET4) treatment from MET4-IO participants (Fig. 1A). Stools from mouse recipients were sequenced alongside human donor stool to identify differences in stool composition between donor-recipient pairs. Human donor stools were represented by a large proportion of Bacillota with some Actinomycetota and Bacteroidota, while mouse recipient stools were dominated by Bacteroidota, Bacillota, and Verrucomicrobiota (Supp Fig. 1). Stool microbiomes of HMA mice were more compositionally similar to mice from other donors than the stool microbiome of their human donor. (PERMANOVA, F=5.35, R^2^=0.2, Fig 1B-C, Supp Fig 2). Dispersion differed between human and mouse groups, indicating lower variance in mouse recipient microbiomes compared to human donors (PERMDISP, F=9.31, *p*=0.0058).

**Figure 1.**
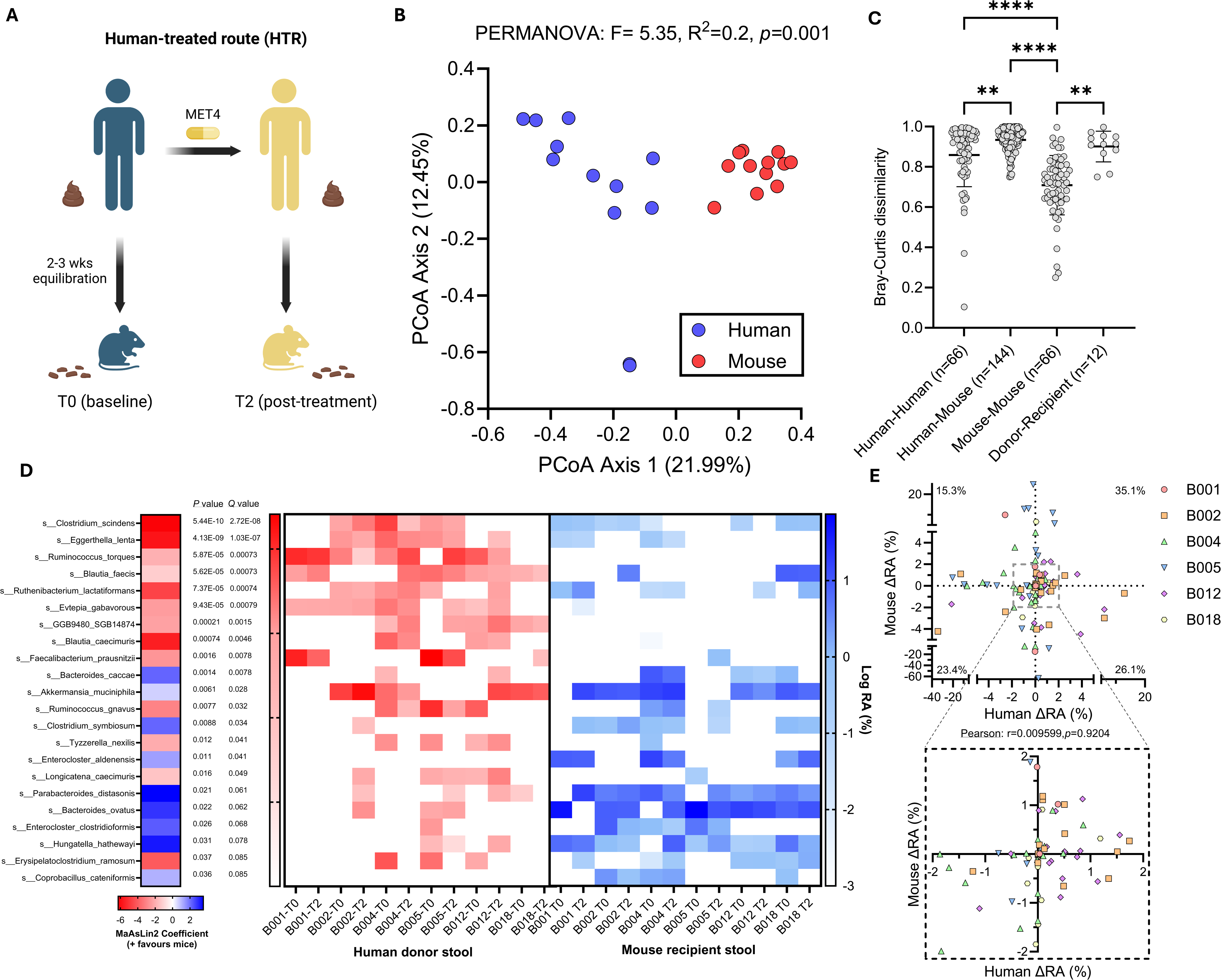
Different gut microbiota engraft in mouse recipients compared to human donors. (A) Flowchart depicting generation of HMA mice. (B) Principal coordinate analysis plot for Bray-Curtis dissimilarity between human donor and mouse stool composition. Each dot is an individual human donor sample or an average of mouse recipient samples from the same donor. PERMANOVA compares species composition. (C) Bray-Curtis dissimilarity for all comparisons between donors, mice, and between donor and respective recipients. Lines indicate the mean and whiskers are standard deviation. Kruskal-Wallis test followed by Dunn’s multiple comparisons test performed. (D) Association between species and relative abundance of stool taxa in human donors and recipient mice, including donor ID and treatment timepoint as random effects. Blue shading indicates taxa enriched in mice, while red indicates taxa enriched in humans. Statistics and coefficients determined with MaAsLin2. Heatmap displays the log % relative abundance of corresponding taxa in individual human and mouse samples. Zero values are arbitrarily set to −3. (E) Scatter plot of delta % relative abundance (RA) for stool taxa from baseline to post-MET4 timepoints in human donors and mouse recipients. Each dot represents an individual taxon, and only taxa overlapping between humans and mice were included. Percent of overlapping taxa is shown for each quadrant. Pearson’s correlation test was performed.

**Figure 2.**
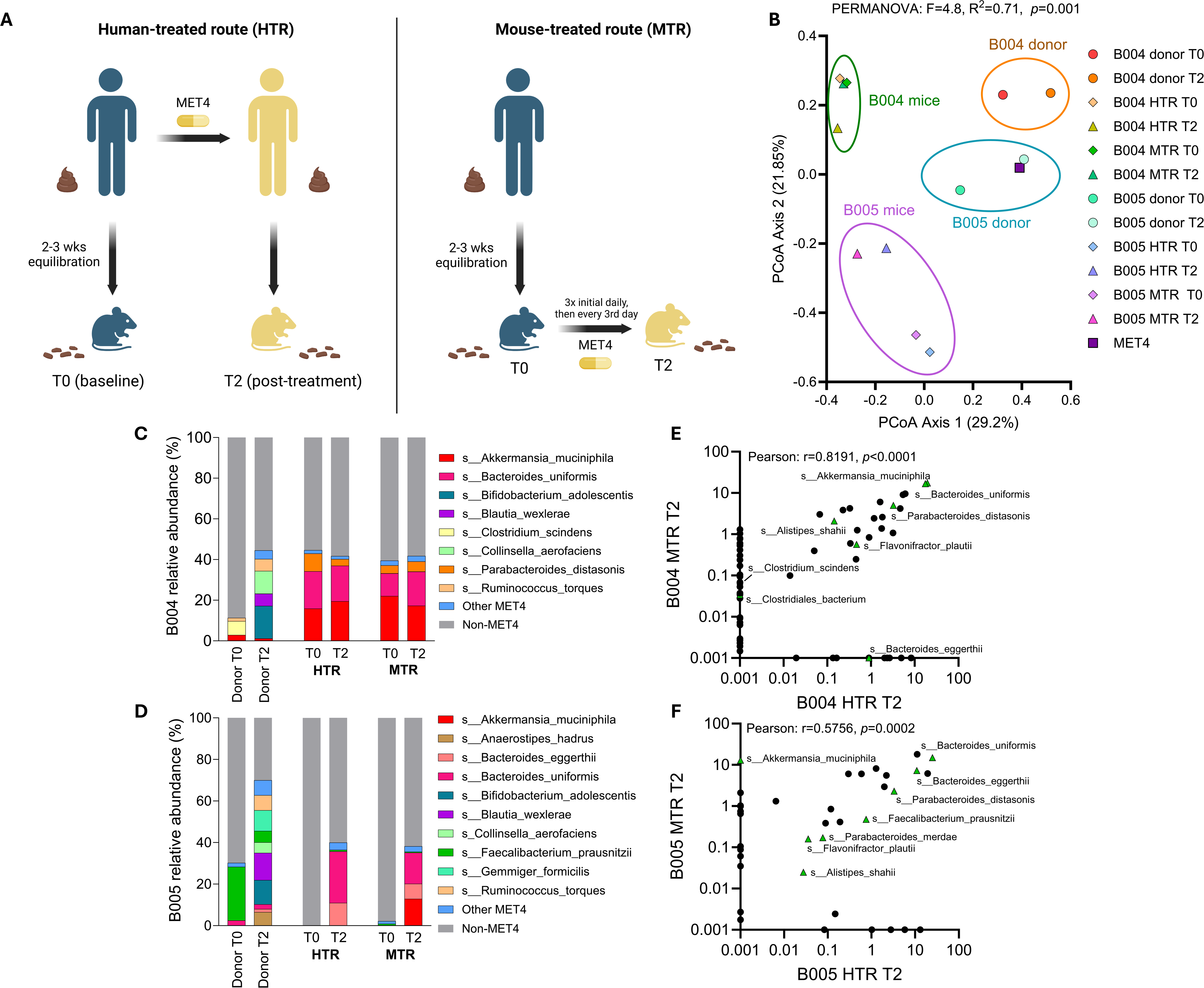
Analysis of the gut microbiome following different treatment exposure routes in germ-free mice. (A) Diagram describing the two routes of treatment exposure in this study. Stools were collected at the indicated timepoints. (B) Principal coordinate analysis plot (PCoA) of Bray-Curtis dissimilarity for MET4, donor stools, and stool from mouse recipients receiving donor FMT and/or direct MET4 administration. Each dot represents an individual human donor sample, an average of MET4 samples sequenced, or an average of mouse recipient stools from the same donor and treatment route. Diamonds represent T0 timepoints, triangles represent T2, and square represents MET4. PERMANOVA compares MET4, donor stool, and mouse recipient stool composition. (C-D) Average relative abundance (%) histograms for stools from mice receiving post-MET4 (T2) donor stool from B004 (D) and B005 (E) or MET4 directly. Donor stool composition is included as a comparator. Non-MET4 taxa are in grayscale and unlabeled. “Other” consists of taxa with <5% relative abundance in every sample. (E-F) Scatter plots showing logscale % relative abundance of gut taxa in B004 (F) and B005 (G) mice compared between routes of exposure at T2. Each point represents a different taxon. MET4 taxa are represented by green triangles.

When controlling for donor and timepoint, specific taxa were found to be enriched in either mouse or human stools, suggesting engraftment bias of distinct taxa depending on host species (Fig. 1D). *Clostridium scindens*, *Eggerthella lenta*, and *Ruminococccus torques* were enriched in human donors, while *Bacteroides caccae*, *Akkermansia muciniphila*, and *Clostridium symbiosum* were increased in mouse recipients (Fig. 1D). T0/T2 differences in % relative abundance of overlapping taxa between donor-recipient pairs revealed poor correlativity, indicating taxonomic changes after treatment in humans were not recapitulated in corresponding HMA mice (Fig. 1E). Only a subset of taxa in donor stools appeared in the stool of recipient mice, with variable engraftment (5.13%-34.69%) across donors (Supp. Fig 3).

### HMA mice display donor-dependent differences in ecology after treatment exposure

Changes in microbiome composition in HMA mice receiving MET4-treated human (T2) stool include all aggregate exposures experienced by the human donor (including diet, medical interventions, and other factors) and may not be solely attributable to consortium exposure. We therefore evaluated ecological changes after direct inoculation of MET4 to assess ecological effects in the absence of these potential confounders. We administered MET4 directly to HMA mice (“mouse-treated route”, MTR) after prior gavage with T0 donor stool to compare taxonomic stool composition to mice receiving T2 donor stool (“human-treated route”, HTR, Fig. 2A). We generated these mice with stool from two donors (B004 and B005) in which a significant proportion of MET4 engrafted (Supp. Fig. 4A).

To assess differences in stool composition between MTR and HTR routes, Bray-Curtis dissimilarity was calculated between mouse samples, donor stools, and the MET4 consortium itself. Distinct compositional clusters were observed for human donors and mouse recipients that were donor-specific (PERMANOVA, F=4.8, R^2^=0.71, Fig. 2B). For each set of mice generated from the same donor, mouse microbial communities were compositionally similar regardless of the route of MET4 exposure. When comparing Bray-Curtis dissimilarity, MET4 and donor stools were highly dissimilar to MTR stools, while MET4-exposed mouse stools were more similar regardless of exposure route (Supp. Fig. 4B). Mice receiving B004 donor stool were more compositionally similar to one another than those receiving B005 donor stool (Supp. Fig. 4C-D) regardless of method by which they were generated (Supp. Fig. 4C, E), indicating changes in HMA mouse microbial ecology after treatment depended on the donor.

### Compositional similarities in HMA mice generated after therapeutic microbial consortium exposure are driven by engraftment of a limited set of taxa

To identify donor-dependent differences in ecology, we compared the percent relative abundance of MET4 taxa between donor stools and routes of treatment exposure for each group of recipient mice. Mice receiving stool from B005, but not B004, displayed increases in MET4 relative abundance by both exposure routes (Fig. 2C-D, Supp. Fig. 5A-B). The dominant engrafting consortium microbes in B005 recipient mice resembled those in B004 mice, which displayed engraftment of MET4 taxa at T0, indicating presence of these taxa even without MET4 exposure in B004 mice. There was a positive correlation in the relative abundance of taxa present in HMA mice regardless of MET4 exposure route (Fig. 2E-F, Supp. Fig. 5C-D). *Alistipes shahii*, *Parabacteroides distasonis*, *Bacteroides uniformis*, *Bacteroides eggerthii*, and *Flavonifractor plautii* were present in post-treatment mice regardless of donor stool or treatment exposure route, suggesting that engraftment of MET4 microbes may be taxonomically restricted in HMA mice.

### Engraftment of a restricted set of taxa is reproducible in HMA mice generated from donors with a variety of health states

To assess whether taxonomically restricted engraftment of donor microbes in HMA mice was unique to our experiments, we analyzed datasets from four studies in which HMA mice were generated from phenotypically diverse human health states, including colon cancer, autism spectrum disorder, inflammatory bowel disease and Parkinson’s disease (Table 1,(20–23)). Stool microbiome composition from human donors and mouse recipients was compared after processing and analyzing publicly available sequencing data from these studies through a single analytical pipeline.

As we observed in our experiments, microbiota in HMA mice generated from diverse populations in unrelated studies were more alike to one another than they were to the human donors from which they were derived (Fig. 3A-D). Correlation of taxonomic relative abundances between human donors and mouse recipients varied by study (range: 0.439-0.969, Fig. 3A). However, the degree of agreement between mouse-mouse pairs (mean: 0.935) was higher than for human-mouse pairs (mean: 0.634) and human-human pairs (mean: 0.580), even when mice were derived from donors from different studies (range: 0.901-0.957, untransformed coefficients shown here, Fig 3B, Supp. Fig. 6).

**Figure 3.**
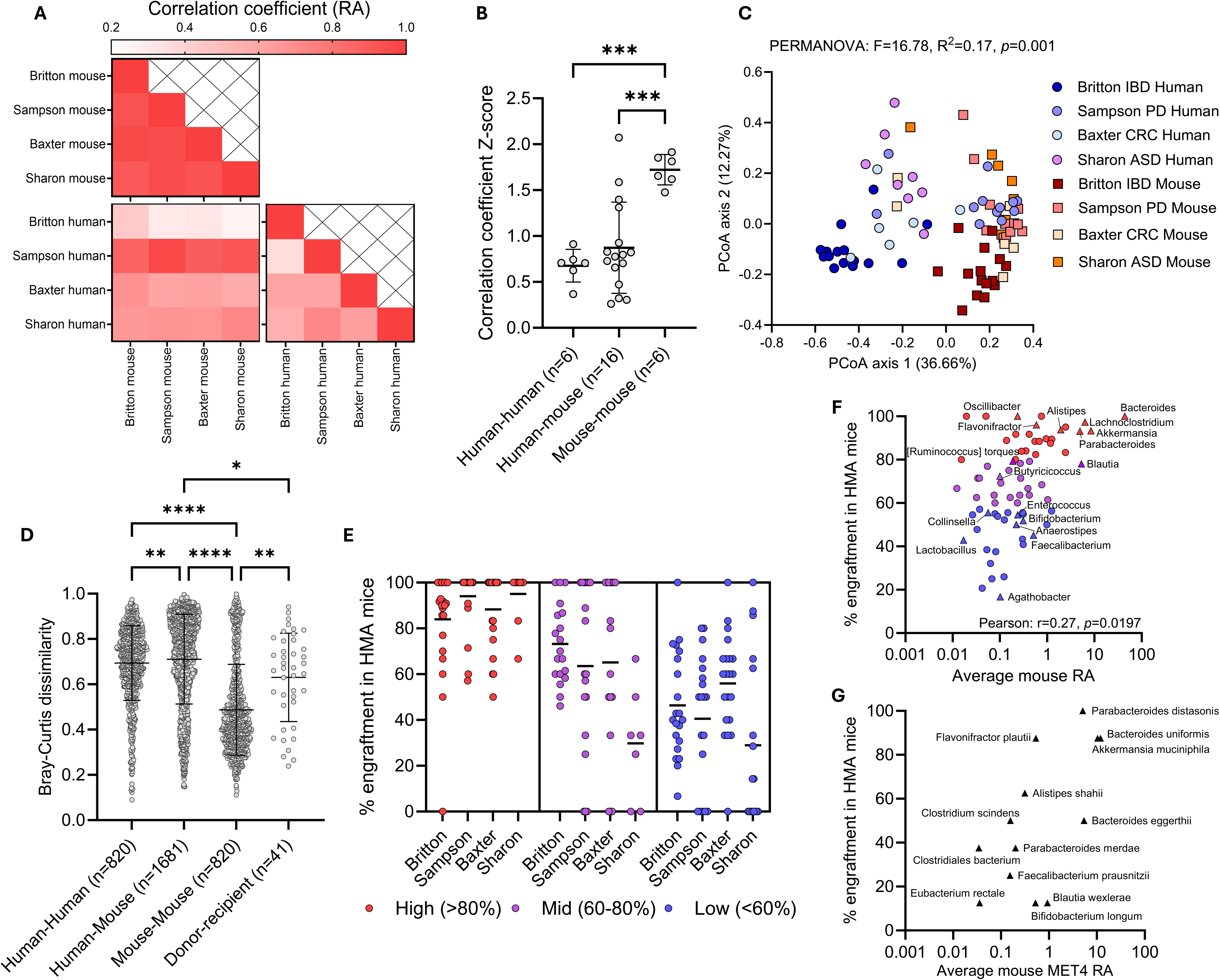
HMA mice engraft specific taxa more consistently. (A) Heat map displaying correlation coefficient for relative abundance (RA) of all taxa between studies and species. (B) Comparison of Z-transformed correlation coefficients by ANOVA followed by Tukey’s multiple comparisons test. (C) Principal coordinate analysis plot (PCoA) of Bray Curtis dissimilarity comparing HMA mice to human donors from different studies. Each dot represents a human donor or an average of mouse recipient stools. Circles represent humans, squares represent mice. PERMANOVA compares species composition. (D) Bray-Curtis dissimilarity for all comparisons between donors, mice, and between donor and respective recipients. Kruskal-Wallis test followed by Dunn’s multiple comparisons test performed. (E) Percent (%) engraftment of taxa from HMA mouse study human donors into recipient mice. Only taxa engrafting in >4 mice are included. Each dot represents % engraftment of one taxon in individual studies and lines represent the mean across taxa within the study. Taxa were stratified into high, medium, or low engrafters based on average % engraftment across studies. (F-G) Scatter plots of % engraftment against average mouse RA across HMA mouse studies (F) or average mouse RA of MET4 taxa in mice exposed to MET4 (G). Each dot represents an individual taxon. MET4 taxa are labeled and are represented by triangles. Coloured taxa in (F) correspond to high, medium, or low engraftment from (E). Lines represent mean and whiskers represent the standard deviation in all figures. IBD – intestinal bowel disease; PD – Parkinson’s disease; CRC – colorectal cancer; ASD – autism spectrum disorder

Bray-Curtis dissimilarity analysis between human donors and mouse recipients from each study revealed distinct clustering of human stools from mouse stools across cohorts (PERMANOVA, F=16.78, R^2^=0.17, Fig. 3C) and differences in dispersion between species (PERMDISP, F=31.63, *p*<0.0001), indicating lower variance in HMA mouse gut microbiomes as we observed in our cohort. Comparison of Bray-Curtis dissimilarities between species presented greater similarity in HMA mice across studies compared to the human donors they were generated from (Fig. 3D).

Notably, one human cohort (Sampson, Parkinson’s disease) displayed strong correlations with cross-study mouse relative abundances (Fig. 3A) and clustered closely with HMA mouse microbiomes (Fig. 3C), suggesting this high degree of similarity was from a human cohort with stool samples that already resembled HMA mice ecology.

To assess whether engraftment was a feature of the taxon rather than the experiment, taxa in each study were stratified into high (>80%), medium (60-80%), or low (<60%) engrafters based on their average percent engraftment across HMA mouse studies. ‘High’ engrafters demonstrated concordance across studies (i.e., were high engrafters in all studies), with moderate and low engrafters demonstrating greater heterogeneity within and between studies (Fig. 3E). Within the high engraftment taxa, *Bacteroides* was the most abundant and consistent engrafter amongst all microbes, with all donors transferring this taxon into their corresponding recipient mice when it was present in the donor (Fig. 3F, Supp. Table 2). Among MET4 taxa, *Lachnoclostridium*, *Flavonifractor*, *Alistipes*, *Akkermansia*, and *Parabacteroides* engrafted into mice in over 85% of donor-recipient pairs, while *Collinsella*, *Enterococcus*, *Bifidobacterium*, *Anaerostipes*, *Faecalibacterium*, and *Lactobacillus* engrafted in less than 60% of donor-recipient pairs (Fig. 3F, Supp. Table 2). Percent engraftment into HMA mice was positively correlated with average relative abundance in these mice, suggesting taxa that frequently engraft may also dominate the gut microbiome of mouse recipients (Fig. 3F).

Notably, *Parabacteroides distasonis*, *Bacteroides uniformis*, *Flavonifractor plautii*, and *Akkermansia muciniphila* were the most consistent MET4 engrafters in our cohort as well, being present in over 80% of mice exposed to MET4 via donor stool or MET4 directly (Fig. 3G, Supp. Table 3). Furthermore, *Alistipes shahii*, *Bacteroides eggerthii*, and *Clostridium scindens* were present in over 50% of mice post-MET4 exposure. Of the 13 MET4 taxa with engraftment in our HMA mice, 8/1/3 were high/mid/low engrafters in published studies (Supp. Table 3), while 1 taxon was not clearly identifiable based on annotation.

## Discussion

It has been well established that differences in human and mouse gut ecology exist due to the physiological and environmental differences present between species(1, 3). HMA mice are thought to circumvent these differences with a “humanized” gut microbiome, allowing their use as avatars of human gut ecology. In this study, we sought to assess the ecological fidelity of HMA mice as avatars of humans in an interventional trial of a microbiome-targeting therapeutic. Stool microbiome composition differed between human donors and mouse recipients but was similar across mice irrespective of human donor. A limited repertoire of microbes engrafted into HMA mice, consisting primarily of *Bacteroides*, *Akkermansia*, and *Parabacteroides* species. These results were corroborated in a mega-analysis of four published HMA mouse studies of diverse human health states.

The similarity in composition between HMA mice across studies and the propensity for specific microbes to engraft in these mice suggests that the intrinsic differences between species may overwhelm or mask the ecological effects of disease or treatment phenotypes that studies attempt to causally assess. As an example, in our mice, any changes induced by MET4 treatment in the B004 donor were not observable in recipient mice due to the presence of engraftment-favoured MET4 microbes (e.g. *Bacteroides*, *Parabacteroides*, *Akkermansia*) at baseline (T0) and the absence of several microbes that increased post-treatment in the original donor. Interestingly, strong HMA mouse engrafters, identified both in our cohort and in published datasets, are known to be common colonizers of the mouse colon and associated mucosa(28), suggesting these mice may re-create typical mouse gut ecology based on what is present in the inoculum.

Additionally, across published HMA mouse studies, greater engraftment frequency was correlated with greater relative abundance, indicating that strong engrafters were both highly prevalent across HMA mice and highly abundant in the recipient gut. Dominance of these microbes may lead to ecological exclusion of weaker engrafters through niche/resource competition(29), or poorly engrafting taxa may simply be maladapted to the mouse gut and/or diet. Altogether, these findings suggest HMA mice are limited avatars of human gut ecology and therefore may not be appropriate models for evaluating human microbiome-targeting therapeutics, such as complex microbial consortia like MET4.

Our study has important technical and theoretical limitations. Although we used relatively common strategies for generating HMA mice, it involved freezing samples after collection instead of fresh processing into fecal slurries, which is likely to selectively affect taxonomic viability (30). In addition, differences in mouse housing environment and operating procedures can lead to differences in microbial composition and engraftment that may be overcome by tuning environmental and dietary conditions(4). We did not explore modifications to experimental conditions that could enhance the ecological fidelity of HMA models. Factors including differences in human and mouse diet, dissimilarity in host species physiology or immunity, dependencies on other human-adapted taxa not usually present in the mouse gut, or variability in inoculum viability through processing and in the mouse gastrointestinal tract are potentially modifiers of engraftment(1). Strategies to augment the engraftment of human taxa to improve ecological fidelity could include co-administration of modified humanized diets (31), drugs (e.g. antibiotics), or humanization of mouse immunity (32). We did not aim to exhaust the possible explanations for this taxonomic restriction, but merely to characterize it in commonly used experimental conditions, in particular in the context of interventional studies of therapeutic consortia. Despite these limitations, we observed similarities in composition between our experiments and published HMA mouse studies, suggesting that our observations were mostly generalizable to other HMA studies/models.

Our results highlight the poor ecological fidelity of HMA mice as models of microbial consortium exposure and suggest that host restriction of taxonomic engraftment is a deterministic process. This limits the utility of these mouse models for attributing causality to the changes induced by microbial consortia when administered to humans. While HMA mice remain essential models for mechanistic and reductionist experiments, the reproducibility of taxonomic restriction we observed are important considerations when designing experiments to assess broad ecological effects, for which HMA mice as they are commonly generated may be inappropriate models.

## Declarations

### Ethics approval and consent to participate

Approved by University Health Network Review Ethics Board #18-5950. All participants provided written informed consent.

### Availability of data and material

Available upon reasonable request. Metagenomic sequencing data are available in SRA/NCBI under accession number PRJNA1219628. Mega-analysis sequencing data are available from their respective sources.

## Authors’ contributions

MKW, EA, BC wrote the manuscript. MKW, EA performed analysis and generated figures. LLS, AS, BC designed the MET4-IO study. MKW, AAH, PHHS conducted the study or processed samples. KC, KS, EA-V designed and provided the MET4 consortium. All authors reviewed the manuscript and provided feedback.

## Acknowledgements

We would like to thank Susy Hota, Susan Poutanen, and the Microbiota Therapeutics Outcomes Program for their mouse FMT preparation protocol. Some figures were created in BioRender (https://BioRender.com/m52b649).

---

**Supplementary Figure 1. Relative abundance histograms for human donor stools and mice receiving donor baseline and post-treatment stools.** Relative abundance (%) of phyla for human donor stool and mouse recipient stool samples. Bars from mouse samples represent the average of 2-4 mice receiving stool from the same donor.

**Supplementary Figure 2. PCoA of Bray-Curtis dissimilarity with stratification by human donor.** Principal coordinate analysis plot stratifying samples based on donor. Each dot represents an individual human donor sample or an average of mouse recipient samples from the same donor. Human donor samples are circled in red and mouse recipient samples are circled in blue.

**Supplementary Figure 3. Number of engrafting taxa in mice varies by stool and donor.** Scatter plots depicting logscale % relative abundance of taxa in germ-free mice receiving participant stools from before (T0) and after MET4 treatment (T2). Percent of taxa engrafted from donor is displayed in bottom right of each plot.

**Supplementary Figure 4. Bray-Curtis dissimilarity between routes of MET4 exposure and inoculum.** (A) Log fold change of relative abundance from T0 to T2 for MET4 taxa in donor stools. Areas outside of grey box show taxa with >1 log fold increase or decrease post-treatment. One sample t-tests were performed to assess non-zero fold change. (B) Comparison of Bray-Curtis dissimilarity between MET4, donor stools, and HTR/MTR mouse stools compared to MTR mouse stools. (C) Bray-Curtis dissimilarity heat map between different routes of exposure for B004 and B005 at T0 and T2 timepoints. (D-E) Comparisons of Bray-Curtis dissimilarity between B004 and B005 mice (D) and between intra- and inter-donor routes (E). Each dot in (B, D-E) represents the Bray-Curtis dissimilarity between mice generated from a single donor stool for each inter-route/timepoint/donor comparison. Lines represent mean and whiskers represent the standard deviation in all figures. Kruskall-Wallis followed by Dunn’s multiple comparison test performed for (B, D-E).

**Supplementary Figure 5. Full relative abundance histograms for mice receiving donor FMT and MET4 and scatter plots for routes at baseline T0.** (A-B) Average relative abundance (%) of stool for B004 (A) and B005 (B) from baseline (T0) and post-MET4 (T2) timepoints for human-treated route (HTR) and mouse-treated route (MTR) mice. Non-MET4 taxa are shown in grayscale. “Other” consists of taxa with <5% relative abundance in every sample. (C-D) Scatter plots showing logscale % relative abundance of gut taxa in B004 (C) and B005 (D) mice compared between routes of exposure at baseline T0. Each point represents a different taxon. MET4 taxa are represented by green triangles.

**Supplementary Figure 6. Correlation scatter plots between human and mouse relative abundance in mega-analysis studies.** Scatter plots depicting logscale average % relative abundance of taxa in HMA mice compared to their respective donors. Each dot represents an individual taxon. Pearson correlation performed for each donor-recipient comparison; all comparisons were *p*<0.0001.

**Supplementary Table 1. MET4 taxonomic annotations**

See Excel sheet. Mega-analysis annotations only include taxa that reached the cut-off threshold (>4 total mice engrafted)

**Supplementary Table 2. Commonly engrafting taxa in HMA mouse studies.**

See Excel sheet. Cut-off for inclusion in analysis was >4 total mice engrafted across studies.

**Supplementary Table 3. Engrafting MET4 taxa in consortium-exposed HMA mice.**

See Excel sheet.

